# Genomics-accelerated discovery of diverse fungicidal bacteria

**DOI:** 10.1101/2020.10.16.343004

**Authors:** Matthew B Biggs, Kelly Craig, Esther Gachango, David Ingham, Mathias Twizeyimana

## Abstract

Sorghum Anthracnose and Black Sigatoka of bananas are problematic fungal diseases worldwide, with a particularly devastating impact on small-holder farmers in Sub-Saharan Africa. We screened a total of 1,227 bacterial isolates for antifungal activity against these pathogens using detached-leaf methods and identified 72 isolates with robust activity against one or both of these pathogens. These bacterial isolates represent a diverse set of five phyla, 14 genera and 22 species, including taxa for which this is the first observation of fungal disease suppression. We identified biosynthetic gene clusters associated with activity against each pathogen. Through a machine learning workflow we discovered additional active isolates, including an isolate from a genus that had not been included in previous screening or model training. Machine-learning improved the discovery rate of our screen by 3-fold. This work highlights the wealth of biocontrol mechanisms available in the microbial world for management of fungal pathogens, generates opportunities for future characterization of novel fungicidal mechanisms, and provides a set of genomic features and models for discovering additional bacterial isolates with activity against these two pathogens.

## Introduction

Sorghum (*Sorghum bicolor* (L.) Moench) is one of the top five cereal crops in the world agricultural economy, and second only to maize as a staple for food-insecure people in Sub-Saharan Africa (1). Production of sorghum in Africa is constrained by both abiotic and biotic factors. The major biotic constraint is the fungal disease sorghum anthracnose (SA), caused by *Colletotrichum sublineolum* P. Henn. Kabat et Bub. Many sorghum landraces and improved varieties are susceptible to the foliar stage of the disease and yield losses ranging from 41% to 67% have been reported in Africa (2).

Bananas and plantains (Musa spp.) are also of critical importance to food security and income generation for over 100 million people in Sub-Saharan Africa (3,4). As they produce fruits year round, these perennial plants are the backbone of many farming systems. They also protect the soil from erosion and are resistant to floods and drought. Black Sigatoka disease (BS), also known as black leaf streak, caused by the fungal species *Mycosphaerella fijiensis*, is regarded as the most economically important leaf disease that affects bananas and plantains worldwide (5). The disease results in heavy losses and, in highly susceptible varieties, can lead to the total collapse of the plant.

Bacteria provide a rich resource for the discovery of novel tools to manage plant diseases. Bacteria have co-inhabited many environments with fungal competitors for millions of years and have evolved mechanisms to antagonize and exclude a diverse range of fungal pathogens (6,7). To identify bacteria with these naturally-occurring antifungal activities, we recently completed a large screen of more than 1,227 bacterial isolates against SA and BS. We discovered 72 bacterial isolates that significantly and reproducibly reduced infection of leaf tissue (53 isolates in four phyla that controlled SA in sorghum leaf tissue, and 31 isolates in four phyla that controlled BS in banana leaf tissue). These novel isolates provide potential new biological tools for managing the destructive effects of these pathogens.

Our search for the best fungicidal candidate isolates was accelerated using genomics and machine learning to strategically select isolates from a large microbial culture collection. Through this extensive screening program, we used two distinct methods to identify genomic features (Biosynthetic Gene Cluster families or “BGC families”) that are predictive of bacterial activity against SA and BS. These features suggest an expanded pathogen spectrum for some known fungicidal molecules, include entirely novel molecules with unknown modes of action (MOA), and provide a foundation for further research. We provide representative sequences for the most predictive BGC families.

Finally, to demonstrate the predictive value of the BGC families discovered in this work, we used machine learning models. We evaluated the strengths and limitations of multiple modeling frameworks *in silico* (including neural networks, random forests, and feature enrichment). We found that random forest models trained on oversampled data performed the best, followed closely by deep neural networks. In conjunction with these computational evaluations, we selected 176 additional bacterial isolates using two of the machine learning methods and screened them for antifungal activities. This additional round of model-driven screening led to the identification of a new genus (which had not been included in previous screening or model training) with activity against both pathogens and a 3-fold increase in our screening discovery rate. We provide the resulting machine learning models as tools for the community.

## Materials and Methods

### Microbial isolation and genome sequencing

Samples were collected from various soil samples, leaves, roots, insects, plants, amphibians, tubers, grain bin dust, and excrement. Samples were collected in the United States and Africa from 2013 to 2019 (Figure 1A). Rhizosphere samples were collected by washing soil attached to the roots with 0.1M sodium phosphate buffer (pH7) and filtering through a 40 μm filter. Insects and plant parts were ground with a mortar and pestle, mixed into a slurry with 0.1M sodium phosphate buffer (pH7) and filtered through a 40 μm filter. Environmental samples were plated on oxoid brilliance bacillus cereus agar (Thermo Fisher Scientific), luria-bertani agar (Thermo Fisher Scientific), or modified M9 minimal salts agar. Modified M9 salts consists of, per liter, 11.33g Na_2_HPO_4_7H_2_O, 3g KH_2_PO_4_, 1g NH_4_Cl, 10g C_5_H_8_NNaO_4_H_2_O, 30g Molasses (Grandma’s Unsulphured Molasses), 2mM MgSO_4_7H_2_O, 0.2mM ZnSO_4_7H_2_O, 0.02mM FeSO_4_7H_2_O. All M9 minimal salts materials were ordered from Sigma-Aldrich unless otherwise stated. Isolates were grown in LB broth and frozen in 15% glycerol stocks. The AgBiome isolate collection served as a resource for this work, and at the time consisted of approximately 70,000 isolates.

**Figure 1.**
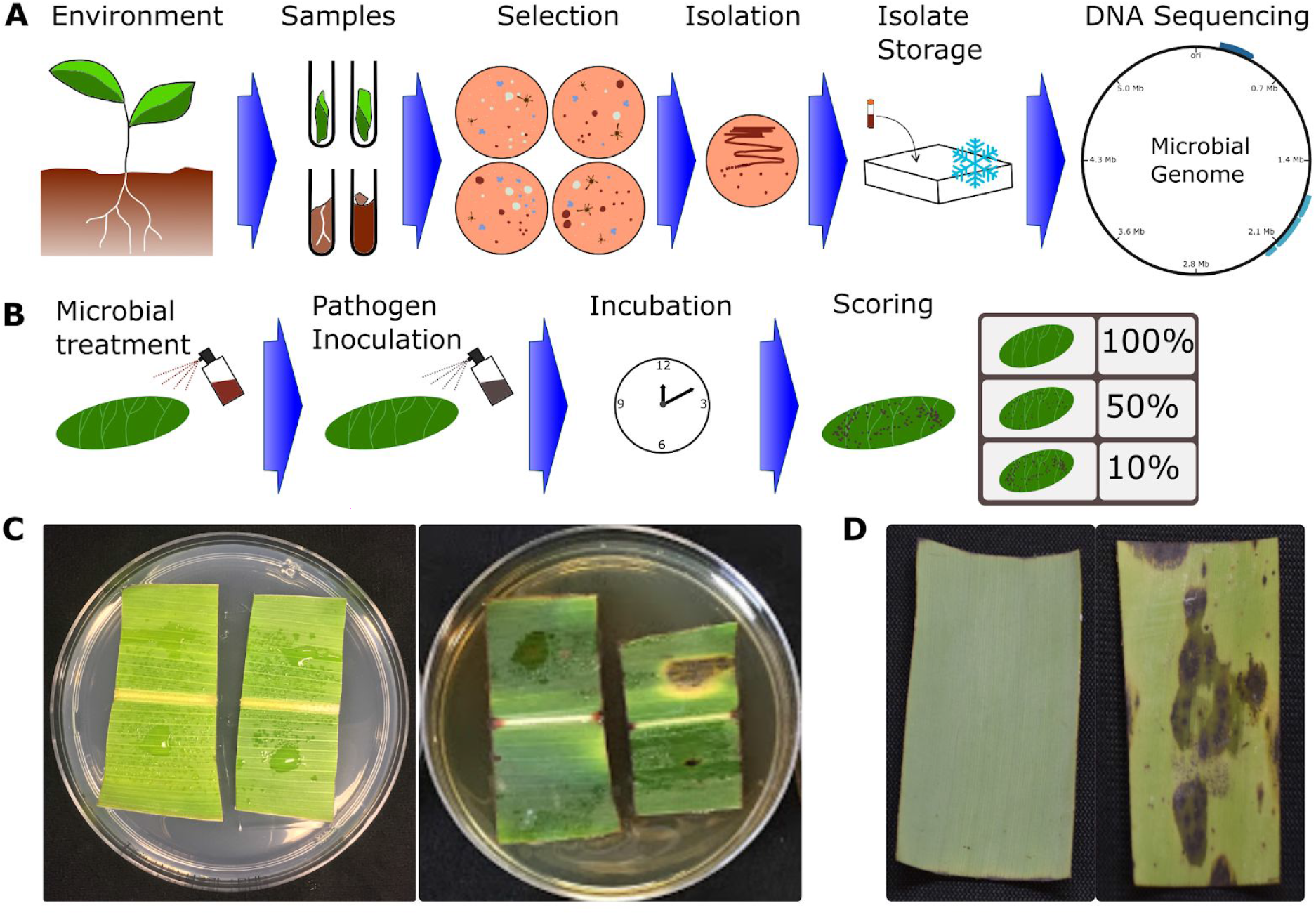
**A.** A high level overview of the bacterial isolation, storage and genome sequencing pipeline. **B.** A high level overview of the leaf disk screens(comparable in principle between SA and BS screens). Leaf disks were embedded in agar, and treated with resuspended bacterial cells. After 24 hours, the leaf disks were treated with fungal spores and incubated for multiple weeks. Scores were assigned by comparing leaf disk disease progression with inoculated and non-inoculated controls. Disease control greater than 70% was considered “active”. **C.** Examples of healthy (left) and diseased (right) sorghum leaf pieces. **D.** Examples of healthy (left) and diseased (right) banana leaf pieces

### Isolate growth and handling

Bacterial isolates were cultured for two days in modified nutrient sporulation medium at 28°C (225 rpm). Modified nutrient sporulation medium consists of, per liter, 5g NaCl (Thermo Fisher Scientific), 10g tryptone (Thermo Fisher Scientific), 8g nutrient broth (BD Biosciences), 0.14mM CaCl_2_ (Sigma Aldrich), 0.2mM MgCl_2_6H_2_O (Sigma Aldrich), and 0.01 mM MnCl_2_4H_2_O (Sigma Aldrich). Bacterial cells were collected and resuspended in 1 mM MgCl_2_ solution.

### Sorghum Anthracnose screen

Sorghum cultivar 12-GS9016-KS585 seeds (supplied by Chromatin Inc.) were grown in the greenhouse for a steady supply of disease-free leaf tissue. Fully expanded sorghum leaves from 4–6 week old plants, were excised, and cut into equal pieces, 2.5 cm wide. To prepare the inoculum, *C. sublineolum* isolated from sorghum in Texas was obtained from Dr. Thomas Isakeit’s laboratory at Texas A&M University, and grown on 20% Oatmeal agar (MilliPORESiGMa, Cat No. 03506), for 14 days. The actively growing fungal culture was flooded with sterile distilled water to dislodge the spores. The concentration of the spore suspension was then adjusted to 1×10^6^ spores/ml. Tween 20 was added to the suspension at the rate of 0.05%.

The bacterial strains were applied to the leaf pieces by spraying 120 μL of the bacterial culture suspended in magnesium chloride buffer (1×10^8^ CFU/ml) using a ribbed skirt fine mist fingertip sprayer (ID-S009, Container & Packaging Supply, Eagle, ID), fitted to a 15 ml conical centrifuge tube (Fisher Scientific, Cat No.14-59-53A). The treated leaf pieces were then plated on 1% water agar amended with 6-Benzylaminopurine (BAP) and incubated at room temperature in the dark (Figure 1B). 24 hours post treatment, the leaf pieces were inoculated with a 30 μL droplet of *C. sublineolum* spore suspension, applied on each side of the mid-rib. The plates were then incubated in a growth chamber (Percival Scientific, Inc) set to a 12-hour photoperiod, maintained at 25°C and 95% relative humidity. The experimental design was a randomized complete block design with 3 replications.

Seven days post inoculation, the leaf pieces were assessed for anthracnose severity on a scale of 0-4 according to (8), with few modifications (Figures 1B and 1C). 0-No symptoms or chlorotic flecks, 1-hypersensitive reaction with no acervuli, 2-lesions with minute and few acervuli, 3-lesions with minute and few acervuli ≤25% of the leaf tissue and 4-lesions with acervuli covering ≥25% of the leaf surface. To identify isolates with robust, reproducible activity, we performed a “confirmation screen” on all active isolates from the primary screen. The confirmation screen was conducted using the same protocol as the primary. Data was analyzed using analysis of variance (ANOVA) in JMP^®^ (version 14.0.0; SAS Institute Inc., Cary, NC).

### Black Sigatoka screen

The susceptible *musa* cultivar Grand Nain (supplied by Green Earth, Melbourne, Florida) was used in this experiment. Plants were maintained in the greenhouse for a constant supply of disease-free leaves. A *Mycosphaerella fijiensis* culture (ITC0489) obtained from the International Institute of Tropical Agriculture (IITA), Ibadan, Nigeria was used for inoculation.

Bacterial isolate application and inoculation were performed as follows: smaller leaf pieces (4 cm long × 3 cm wide) were cut from the excised leaf. Two of these pieces were placed in plastic petri dishes with an adaxial side appressed on water agar amended with 5 mg/liter gibberellic acid (Twizeyimana et al. 2007). Leaf pieces were sprayed with 120 μL of bacterial isolate (1 × 10^8^ CFU/ml of sterile distilled water) using a fingertip sprayer (Container & Packaging Supply, Eagle, ID) fitted to a 15 ml conical centrifuge tube (Fisher Scientific, Cat No.14-59-53A). Petri dishes with leaf pieces were incubated at room temperature in the dark. 24 hours later, leaf pieces in petri dishes were inoculated with a mycelial suspension of *M. fijiensis* (120 μl per leaf piece) using a fingertip sprayer fitted to a 50 ml conical centrifuge tube. The suspension contained mycelial fragments scraped from growing cultures that were cut into smaller mycelial tips in sterile distilled water in 50 ml conical tubes using a homogenizer (Omni International, Kennesaw, GA). The suspension was filtered through two layers of cheesecloth and then stirred. 0.05% Tween 20 and 0.02% Silwet L-77 (Loveland Industries Inc., Greeley, CO) were added. Using a hemocytometer, the suspension was adjusted with sterile distilled water to a concentration of 1 × 10^6^ mycelial fragments/ml. A day after inoculation, plates were incubated in a growth chamber (Percival Scientific, Inc) set at 14 hours photoperiod, maintained at 25°C and 90% relative humidity.

Data recorded were the most progressed stage on inoculated leaves at the time of data collection (9). Data were analyzed using analysis of variance (ANOVA) in PROC GLM of SAS (version 9.4; SAS Institute Inc., Cary, NC) and significant differences (*P* < 0.05) were observed among treatments. To identify isolates with robust, reproducible activity, we performed a “confirmation screen” on all active isolates from the primary screen. The confirmation screen was conducted using the same protocol as the primary.

### Bacterial genomics

Bacterial isolates were grown in liquid culture, spun down, and DNA was isolated from the cell pellets using MoBio microbial DNA isolation kits (Qiagen, Hilden, Germany). The resulting DNA was quantified using a Quant iT PicoGreen assay (Invitrogen, Carlsbad, CA, USA). One nanogram of quantified DNA was sheared enzymatically at 55°C for 5 min using the Illumina Nextera XT tagmentation enzyme (Illumina, San Diego, CA, USA). Tagmented DNA fragments were enriched by 10 cycles of PCR amplification using PCR master mix and primers with the index from Illumina. Libraries were quantified by the KAPA SYBR fast quantitative PCR kit (Life Technologies, Carlsbad, CA, USA) and pooled at a 4 nM concentration. Libraries were denatured with 0.2 N NaOH and sequenced on an Illumina HiSeq sequencing platform. Illumina paired-end reads were demultiplexed using Illumina software bcl2fastq v2.18.0.12. Paired-end reads were adapter and quality trimmed using cutadapt version 1.5 as recommended by Illumina. Trimmed paired-end reads were assembled and reads were aligned back to the consensus sequence using the CLC Genomics programs CLC Assembly Cell and CLC Mapper from Qiagen.

Bacterial genomes were assigned taxonomic identifiers using the Genome Taxonomy Database Toolkit version 1.0.2 (10–12). Biosynthetic gene clusters (BGCs) were annotated using antiSMASH version 5 (13) with the following options enabled: taxon “bacteria”, KnownClusterBlast against MIBiG (14) and General ClusterBlast, Active Site Finder, “pfam2go” mapping, and Prodigal (15) as the gene finding tool. We used BiG-SCAPE with default settings (16) to cluster BGCs into families (the default alignment is “glocal” and the default cutoff for clustering into a family is a distance metric of 0.3).

### Enrichment analysis and machine learning

Isolate feature enrichment and BGC family enrichment were calculated using a hypergeometric distribution. Isolates were considered “active” if they controlled disease progression by 70% or greater. Specifically, we estimated the cumulative probability that given *M* isolates screened, with *n* of the total having a particular attribute (e.g. are derived from soil), and *N* of the total displaying activity, that we would observe *o* or more active isolates with that attribute by chance alone. Smaller probabilities indicate “enriched” attributes that are unlikely to be so highly represented given chance alone, and that potentially have a meaningful biological explanation. We leveraged the hypergeometric module in SciPy to calculate enrichment p-values (17).

We used SciKit-Learn to split training and test sets, train random forest models and generate predictions, perform cross validation, estimate permutation feature importance, and evaluate model performance (18). Permutation feature importance (PFI) is a measure of how model performance is affected by randomly shuffling feature columns. PFI is a more robust measure of feature importance than random forest feature importance (19). Before implementing the random forest models, we tested several configurations of random forest hyperparameters, and selected ensembles of 5 trees with a maximum depth of 32 nodes to maximize recall in a 50-fold cross validation analysis. Oversampling was performed by replicating all the active observations in the training data, thereby increasing the ratio of active to inactive observations. Precision, recall and F1 scores were calculated using the SciKit-Learn functions.

We used PyTorch (20) paired with Skorch (https://github.com/skorch-dev/skorch) to train a deep neural network and generate predictions. We tested several configurations and selected a 3-layer design with 10 fully-connected nodes and reLu activation in the first layer, 30% dropout between the first and second layers during training, 10 fully-connected nodes and reLu activation in the second layer, and a single sigmoid output node to predict the probability of activity. We used a learning rate of 0.02, trained each model for 10 epochs using the Adam optimizer and the binary cross entropy loss function.

To select additional isolates for screening, we selected representative “query” BGCs for each enriched or important BGC family and clustered them with batches of BGCs from the remainder of our collection using BiG-SCAPE with the same parameters. The BGCs that clustered with a query BGC were assigned to the same family as the query.

The data and code developed for this work are publicly available on GitHub: https://github.com/mbi2gs/fungicidal_bact_genomics

## Results

We initially screened 1,051 cultivable bacterial isolates against SA, and 559 of those same isolates against BS. These 1,051 strains were isolated from 27 states in the United States and five districts in Uganda, and were derived from 11 sample types (e.g. soil, plant tissue, insects). We balanced taxonomic breadth and depth by screening more deeply among taxa with well-known fungicidal activity (e.g. *Bacilli* and *Pseudomonads*) while also ensuring that we sampled from the broader diversity of the collection. We refer to this first pool of tested bacterial isolates as “diversity” screening (546 isolates against SA; 145 isolates against BS). In addition, we included isolates that showed activity in other screens against different fungal pathogens, referred to this pool as “spectrum” screening (291 isolates against SA; 259 isolates against BS). Finally, in several cases when an isolate displayed activity against a pathogen, we followed up by screening additional isolates from the collection with high degrees of genomic similarity to that first active isolate. We refer to this pool of isolates as “genomic similarity” screening (214 against SA; 155 against BS).

Viewing the results from a taxonomic perspective, we discovered 156 isolates that displayed primary activity against SA, BS or both, representing five phyla, 27 genera, and 37 species (based on Genome Taxonomy Database labels (11)). We found 72 reproducibly (active again in the confirmation screen) active isolates against SA or BS in five phyla, 14 genera, and 22 species (Figure 2). Isolates that reproducibly controlled both SA and BS came from two phyla, specifically, Proteobacteria and Firmicutes. The active isolate from the phylum Firmicutes_I only controlled SA, the active isolate from the phylum Bacteroidota only controlled BS, and the active isolates from Actinobacteriota only controlled one disease or the other (Figure 2). Of the 95 genera screened against SA, 19 were active in the primary screen and nine showed reproducible, robust control of SA in the confirmation. Of the 44 genera screened against BS, 12 were active in the primary screen, and eight showed reproducible control of BS. Viewing the results from a discovery rate perspective, of the 546 isolates from the diversity screen against SA, 28 controlled SA in the primary (a discovery rate of 5.0%), and eight repeated the control in the confirmation screen (a discovery rate of 1.5%). Of the 145 isolates from the diversity screen against BS, 13 controlled the pathogen in the primary (a discovery rate of 9.0%) and two repeated the control in the confirmation screen (a discovery rate of 1.4%).

**Figure 2.**
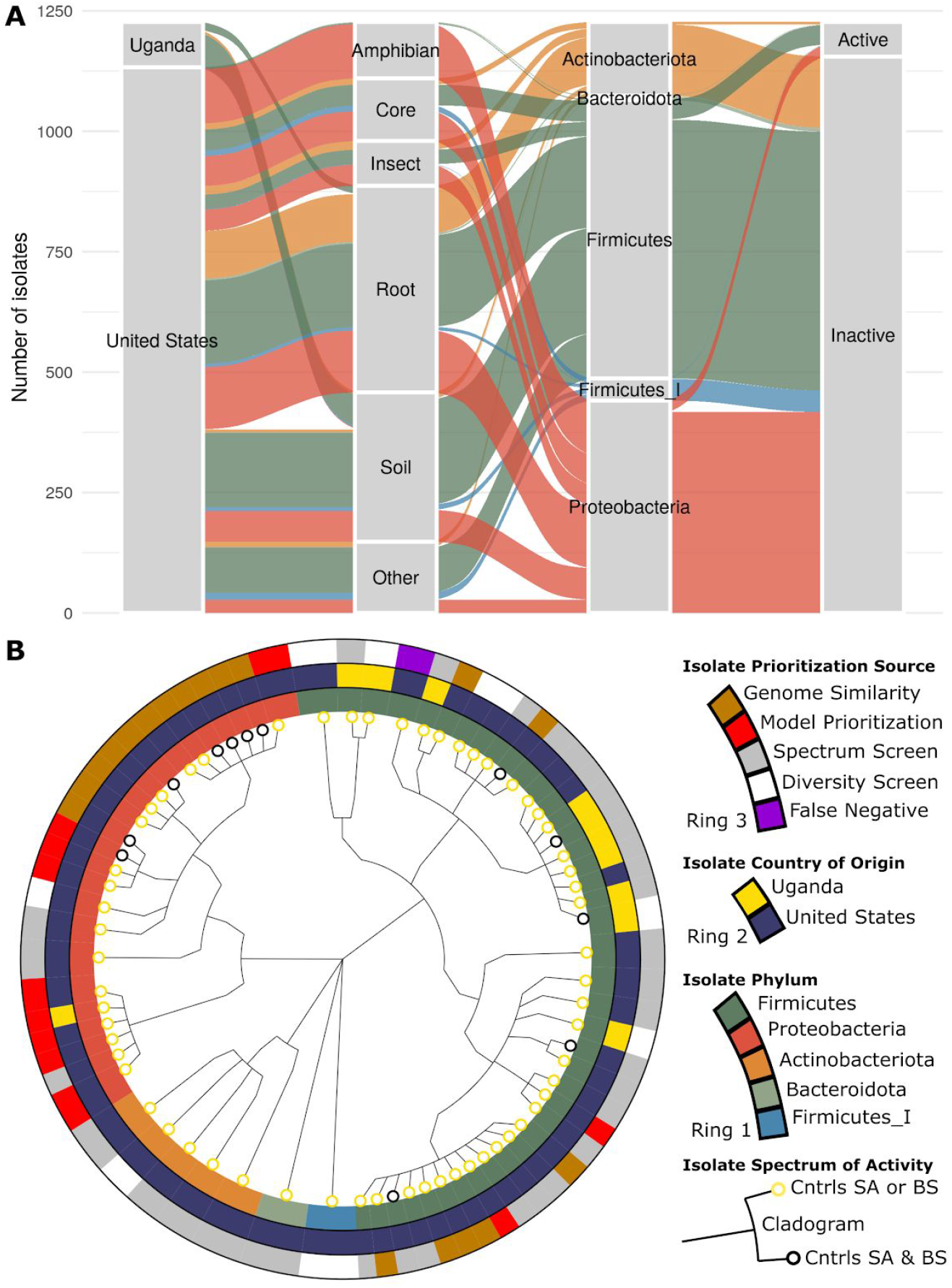
Scope of screening effort, and diverse isolates with confirmed fungicidal activity. **A**. The alluvial diagram illustrates the source country, sample type (the less abundant types were combined into “Other”), phylum membership and fungicidal activity of the 1,227 isolates screened in this work. **B.** The cladogram in the center highlights the diversity of the biocontrol isolates discovered in this work (five phyla represented by the center branches, 14 genera, 22 species). Multiple isolates displayed activity against both diseases (black leaf nodes). The outer ring shows the isolate prioritization strategies that led to each discovery, including predictive models. The “false negative” isolate in purple was predicted to be inactive by all modeling approaches and yet was found to be reproducibly active against SA.

We evaluated the association between fungicidal activity (against SA or BS), and the associated taxonomy, source environment, isolation protocol, and geography. The metric used was an “enrichment score” (see Materials and Methods). After False Discovery Rate (FDR) correction, no p-values were below the chosen threshold of 0.05 (see Table 1). However, the variables with the strongest association were taxonomic, specifically members of the class Bacilli, where the species *Bacillus velezensis* were most enriched for activity against either pathogen, The strongest non-taxonomic associations were with isolates from the soil environment, and isolates from Uganda. Co-variances between each category are presented in Supplemental Figure 2.

**Table 1.**

| Table 1 |  |  |  |  |  |
| --- | --- | --- | --- | --- | --- |
| Variable | Category | Num Actives in Category | Total Strains in Category | P-value | FDR-corrected p-value |
| species | Bacillus velezensis | 2 | 4 | 0.001 | 0.27 |
| family | Bacillaceae | 3 | 19 | 0.003 | 0.27 |
| genus | Bacillus | 3 | 19 | 0.003 | 0.27 |
| class | Bacilli | 6 | 118 | 0.004 | 0.27 |
| phylum | Firmicutes | 6 | 118 | 0.004 | 0.27 |
| sample_type | Soil | 4 | 48 | 0.005 | 0.27 |
| country | Uganda | 3 | 32 | 0.012 | 0.58 |
| species | MA-N2 sp002009585 | 1 | 1 | 0.016 | 0.69 |
| species | Lysinibacillus pakistanensis | 1 | 2 | 0.033 | 1.00 |
| family | Planococcaceae | 2 | 19 | 0.036 | 1.00 |
| order | Bacillales_A | 2 | 19 | 0.036 | 1.00 |
| genus | Paenibacillus_B | 1 | 3 | 0.049 | 1.00 |
| species | Bacillus_A cereus | 1 | 3 | 0.049 | 1.00 |
| species | Paenibacillus_B<br>sp001894745 | 1 | 3 | 0.049 | 1.00 |

To identify biosynthetic pathways associated with control of *Colletotrichum* and *Mycosphaerella*, we generated genome assemblies for all isolates. We annotated the genomes using antiSMASH, which produced multiple BGCs and BGC fragments per genome assembly. We clustered the annotated BGCs and fragments into 2,770 families using BiG-SCAPE (see Materials and Methods). We performed an enrichment analysis to associate isolate activity in the SA primary screen with the presence or absence of each BGC family (Table 2). We found that six of the top 12 BGC families most highly associated with SA control were homologous to well-known fungicidal pathways such as Fengycin (21,22) and Pyrrolnitrin (23,24), validating the enrichment approach for identifying biologically-relevant genomic features. Four of the top 12 families did not show significant homology to well-characterized BGCs (based on the antiSMASH Known Cluster Blast against the MIBiG reference database). These four BGC families are associated with the genus *Bacillus_A*. Three are described as bacteriocins and one as non-ribosomal peptide synthetase-like (NRPS-like) (Table 2). These four BGC families represent potentially novel biocontrol pathways and molecules with relevant activity against SA.

**Table 2.**

|  |
| --- |
| Table 2 |

| BGC Family ID | 99th Percentile Importance | Top Enrichment Scores | Predicted Product Class | Homology to Known BGC | Taxonomic Distribution |
| --- | --- | --- | --- | --- | --- |
| ripps2539 | TRUE | TRUE | bacteriocin | None | genus:Bacillus_A |
| nrps2550 | TRUE | TRUE | NRPS | Elsinochrome A | genus:Bacillus_A |
| others2545 | TRUE | TRUE | betalactone | Fengycin | genus:Bacillus_A |
| pksnrp3708 | TRUE | TRUE | NRPS;T1PKS | Zwittermycin A | genus:Bacillus_A |
| terpene2537 | TRUE | FALSE | terpene | Molybdenum cofactor | genus:Bacillus_A |
| nrps2808 | TRUE | FALSE | NRPS | Thiazostatin | genus:Pseudomonas_E |
| terpene4046 | TRUE | FALSE | terpene | Molybdenum cofactor | genus:Bacillus_A |
| ripps3967 | TRUE | FALSE | bacteriocin | None | genus:Bacillus_A |
| nrps9204 | TRUE | FALSE | NRPS | None | genus:Bacillus_A |
| others2542 | TRUE | FALSE | siderophore | Petrobactin | genus:Bacillus_A |
| ripps2548 | TRUE | FALSE | lanthipeptide | None | genus:Bacillus_A |
| terpene6941 | TRUE | FALSE | terpene | None | species:gtdb_novel_strain |
| nrps4047 | TRUE | FALSE | NRPS;T1PKS | Zwittermycin A | genus:Bacillus_A |
| ripps9214 | TRUE | FALSE | sactipeptide | Thurincin H | species:Bacillus_A<br>thuringiensis_J |
| nrps2540 | TRUE | FALSE | NRPS | Zwittermycin A | species:Bacillus_A<br>thuringiensis_J |
| ripps2935 | TRUE | FALSE | bacteriocin | None | genus:Bacillus_A |
| nrps8318 | TRUE | FALSE | NRPS | Polyoxypeptin | genus:Bacillus_A |
| others3835 | TRUE | FALSE | lanthipeptide | Zwittermycin A | species:Bacillus_A<br>thuringiensis_J |
| nrps2971 | TRUE | FALSE | NRPS | None | species:Pseudomonas_E<br>sp001547895 |
| nrps10661 | TRUE | FALSE | NRPS | None | species:Bacillus subtilis |
| pksother8188 | TRUE | FALSE | T3PKS | None | genus:Bacillus |
| nrps11511 | TRUE | FALSE | NRPS | Surfactin | genus:Bacillus |
| ripps2937 | TRUE | FALSE | bacteriocin | None | species:Bacillus_A<br>thuringiensis_J |
| nrps1086 | TRUE | FALSE | NRPS | Pyoverdine | species:Pseudomonas_E<br>chlororaphis |
| terpene11261 | TRUE | FALSE | terpene | None | genus:Bacillus |
| others11626 | TRUE | FALSE | lanthipeptide | Zwittermycin A | genus:Bacillus_A |
| ripps5084 | TRUE | FALSE | lanthipeptide | None | order:Bacillales |
| ripps7030 | TRUE | FALSE | bacteriocin | None | species:Brevibacillus<br>reuszeri_A |
| nrps4051 | FALSE | TRUE | NRPS-like | None | genus:Bacillus_A |
| ripps3390 | FALSE | TRUE | bacteriocin | None | species:Bacillus_A toyonensis |
| nrps4795 | FALSE | TRUE | NRPS | Pyoverdine | species:Pseudomonas_E protegens |
| ripps4045 | FALSE | TRUE | bacteriocin;LAP | None | genus:Bacillus_A |
| nrps2943 | FALSE | TRUE | NRPS | Lipopeptide 8D1-1/2 | species:Pseudomonas_E protegens |
| others2810 | FALSE | TRUE | other | Pyrrolnitrin | class:Gammaproteobacteria |
| pksother4889 | FALSE | TRUE | PKS-like | Anikasin | species:Pseudomonas_E protegens |
| nrps4050 | FALSE | TRUE | NRPS | Bacillibactin | genus:Bacillus_A |

Among the top 12 BGC families associated with activity against BS, there were four BGC families that overlapped with enriched BGC families against SA, including nrps2943, nrps4795, others2810, and pksother4889 (Supplemental Table 1). There were seven BGC families with homology to well-known fungicidal pathways, including Pyrrolnitrin, and Zwittermycin A (25). There were also two BGC families associated with BS control that did not show significant homology to well-characterized BGCs, and these were distinct from the four novel BGC families associated with SA control. In this case, the two novel BGC families were associated with the family Pseudomonadaceae and the genus *Pseudomonas_E* and were described as a bacteriocin and a N-acetylglutaminylglutamine amide (NAGGN). These BGC families represent potentially novel biocontrol pathways and molecules with relevant activity against BS.

As an additional metric for associating BGC family with pathogen control, we built a random forest classifier to predict isolate activity in each primary screen and calculated the permutation feature importance of each BGC family (see Materials and Methods). There were 28 BGC families in the top 99th percentile for predicting SA control, and 19 for predicting BS control (Table 2 and Supplemental Table 1). Among those associated with SA control, four BGC families were selected by both the feature importance and enrichment methods. 15 of the 28 selected by feature importance were homologous to well-characterized BGCs, many of which are known to be fungicidal or generally involved in microbial competition, including Zwittermycin A (25,26) and the siderophore pyoverdine (27).13 of the 28 did not share significant homology to well-characterized BGCs, and are described as bacteriocins, NRPSs, lanthipeptides and PKSs. We found that many of the BGC families associated with activity against SA were co-occurring throughout the isolates screened (Supplemental Figure 3). For example, we observed clear clusters of BGC families with strong associations with *Bacillus_A*. Among the BGC families associated with controlling BS (Supplemental Table 1), 11 of the 19 were homologous to well-characterized BGCs, including Bacilysin (28). 8 of the 19 were not significantly homologous to well-characterized BGCs, and are categorized as beta lactones, linear azole-containing peptides (LAPs), bacteriocins and NRPSs. We found that many of these BGC families associated with activity against BS were co-occurring throughout the isolates screened (Supplemental Figure 3). For example, we observed clusters of BGC families strongly affiliated with the species *Pseudomonas_E protegens*. These BGC families without significant homology to well-described BGCs represent potentially novel biocontrol pathways and molecules with relevant activity against SA and BS.

To demonstrate the predictive value of the activity-associated BGC families, we developed computational tools that would increase the discovery rate of fungicidal isolates (Figure 3A). We compared an enrichment-based prioritization method “ER” with random forest models (both standard “RF” and trained on over-sampled data “RFOS”; see Materials and Methods), and a deep neural network “NN” (Figure 3A). We selected an oversampling factor of 13 for the RFOS model based on cross-validation (Supplemental Figure 5). We used only the diversity screening isolates data to train the models. During 100-fold cross-validation with 75/25% training/test split of the data, we repeated the process of feature selection described above (i.e. enrichment and permutation feature importance) using only the subset of training data. After feature selection with the RF, the same features were used to train the NN. As a null model, random isolates were predicted to be active at the same discovery rate as the diversity screen (i.e. 5%). All models were trained to predict activity in the primary screen against SA. We found that all four prioritization tools performed better than the null model in terms of the F1 statistic (p-value of 2×10^−7^ by Kruskal–Wallis test, followed by one-sided Mann–Whitney U tests with p-values < 3×10^−4^) (Figure 3B). The ER method was the poorest performer of the four methods tested, while the RFOS and the deep neural network were the best performers. The RFOS average F1 score was significantly better than ER (p-value < 0.05 by one-sided Mann-Whitney U tests), largely due to the RFOS precision scores. The NN method displayed the best recall, increasing the average recall by 6.5-fold compared to the null.

**Figure 3.**
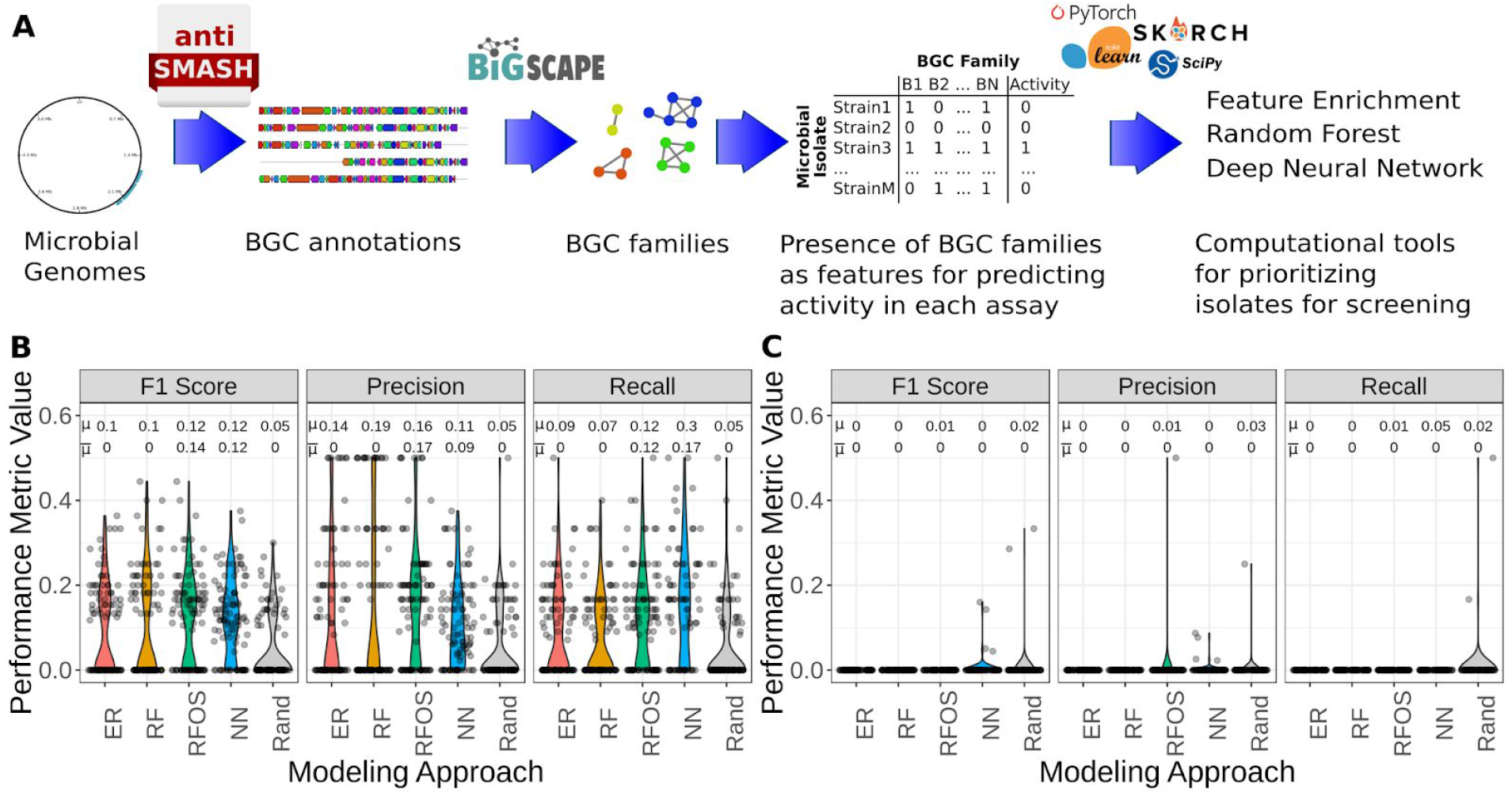
Machine learning workflow and *in silico* results. **A.** Machine learning workflow, starting from genome assemblies, annotating BGCs with antiSMASH, clustering BGCs into families, aggregating the presence or absence of each BGC family in each genome, and finally, selecting features and training models. **B**. Comparing prioritization methods by 100-fold cross validation on the primary screening data. Each iteration of the cross-validation split the data randomly into training and test sets, selected features and trained models on the training data, then predicted activity in the test data. ER (enrichment-based prioritization, where any isolate with one or more enriched BGCs was predicted to be active), RF (random forest model), RFOS (random forest with oversampling), NN (deep neural network), Rand (a random, null model based on the discovery rate from the training data). All prioritization methods outperformed the null model. Both RFOS and NN outperformed the ER and RF models. The mean (μ) and median 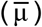 values for each set of simulations are indicated above each distribution. **C.** Each iteration of this cross-validation held out all the members of a particular genus as the test set. In this case, all methods were indistinguishable from the null (Kruskal–Wallis test), indicating that generally speaking, the BGC families defined here are only predictive within a genus. The mean (μ) and median 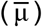 values for each set of simulations are indicated above each distribution.

To determine the extent to which the best performing cross validation iterations were due to taxonomic similarity between the training set and the test set (assuming that more related isolates between the two would improve observed model performance), we compared the Jaccard similarity of the set of genera in the training and test data to the model performance and found no significant correlation (correlation coefficient of 0.097 and a p-value of 0.34 by Spearman’s Rank Correlation; see Supplemental Figure 6). While this result would suggest that taxonomic overlap between training does not affect model performance (which is unlikely), we also repeated the cross-validation evaluation. This time, instead of random training/test splits, we kept out a single genus at each iteration as the test set. We found that under these conditions, the models failed to outperform the null model (p-value 0.13 by Kruskal–Wallis test), suggesting that the BGC families used to train the models tend to be predictive at the genus level or lower (Figure 3C).

In parallel to the *in silico* validation of the predictive value of the enriched BGC families, we also completed an *in vivo* validation. In contrast to the *in silico* validation where we could simulate hundreds of experiments, we performed a single *in vivo* validation. We applied two computational models to predict control of SA from among the larger AgBiome bacterial culture collection. We selected BGC features using enrichment and random forest feature importance, based on an older version of the SA screen data set (the BGC features used in this *in vivo* validation exercise are listed in Supplemental Table 2 and representative sequences are provided in the supplemental materials). To query the collection for isolates with BGCs in the enriched families, we annotated BGCs using antiSMASH across the entire collection, and then used BIG-SCAPE to cluster batches of 200 BGCs together with the queries, using the same clustering parameters as the original clustering job that defined the families. We identified 17,163 isolates that contained at least one representative of the enriched BGC families.

Using two predictive tools (ER and RFOS), we selected 176 isolates from the 17,163 enriched isolates. First, we selected isolates that contained at least one member from the 12 enriched BGC families (ignoring the BGC families identified by permutation feature importance). A large proportion of those isolates belonged to only a few genera, particularly *Bacillus_A*. In order to increase the diversity of the validation set, we required that half of the predicted active isolates come from non-*Bacillus_A* genera. Second, we applied the RFOS model to predict control of SA. We used all fungal-control-associated BGC families as features, trained with 13-fold oversampling, and required that half of the predicted isolates come from non-*Bacillus_A* genera. Finally as a *post hoc* analysis (not to select the 176), we applied the NN model to predict activity of the 176 isolates. We observed that the enrichment method was far more likely to predict that an isolate would be active, and uniquely prioritized 36 isolates, sharing 66 with the RFOS model and 81 with the NN model. The RFOS model prioritized 20 unique solates, and shared 47 with the NN model. The NN model did not prioritize any unique isolates because it was applied *post-hoc*. 20 of the 176 isolates were predicted to be inactive by all three models, one of which was found to be active (this “false negative” isolate was classified as a *Bacillus altitudinis*; see Figure 2).

We found that the RFOS primary discovery rate (precision) was 3-fold greater than the historical discovery rate of the diversity screen (p-value of 3.7×10^−6^ by one-sided two-proportions Z test) (Figure 4). In other words, using the RFOS model, we discovered 3-fold more active isolates for the effort than we would have by screening a random, diverse set of isolates from the collection. The confirmation discovery rate was even higher: 5.4-fold greater than the diversity screen (p-value of 6.2×10^−6^ by one-sided two-proportions Z test) (Figure 4). Additionally, the RFOS model achieved 72% recall, correctly predicting 23 of the 32 total active isolates (p-value of 0.002, compared to 1,000 draws using a random model based on the baseline diversity screen discovery rate of 5.0%) (Supplemental Figure 7). Notably, the RFOS method did not discover any novel taxa that were not already represented in the training set. Most of the true positives were classified as members of the genus *Pseudomonas_E*, while most of the true negatives, false negatives and false positives were classified as members of *Bacillus_A*. Taking into account the SA confirmation screen results (which were not used to train the model), the model precision was not statistically different from the null model, but the recall was exceptionally high, where the RFOS model correctly predicted 11 of the 12 (92%) isolates that reproducibly controlled SA (p-value < 0.001, compared to 1,000 random draws).

**Figure 4.**
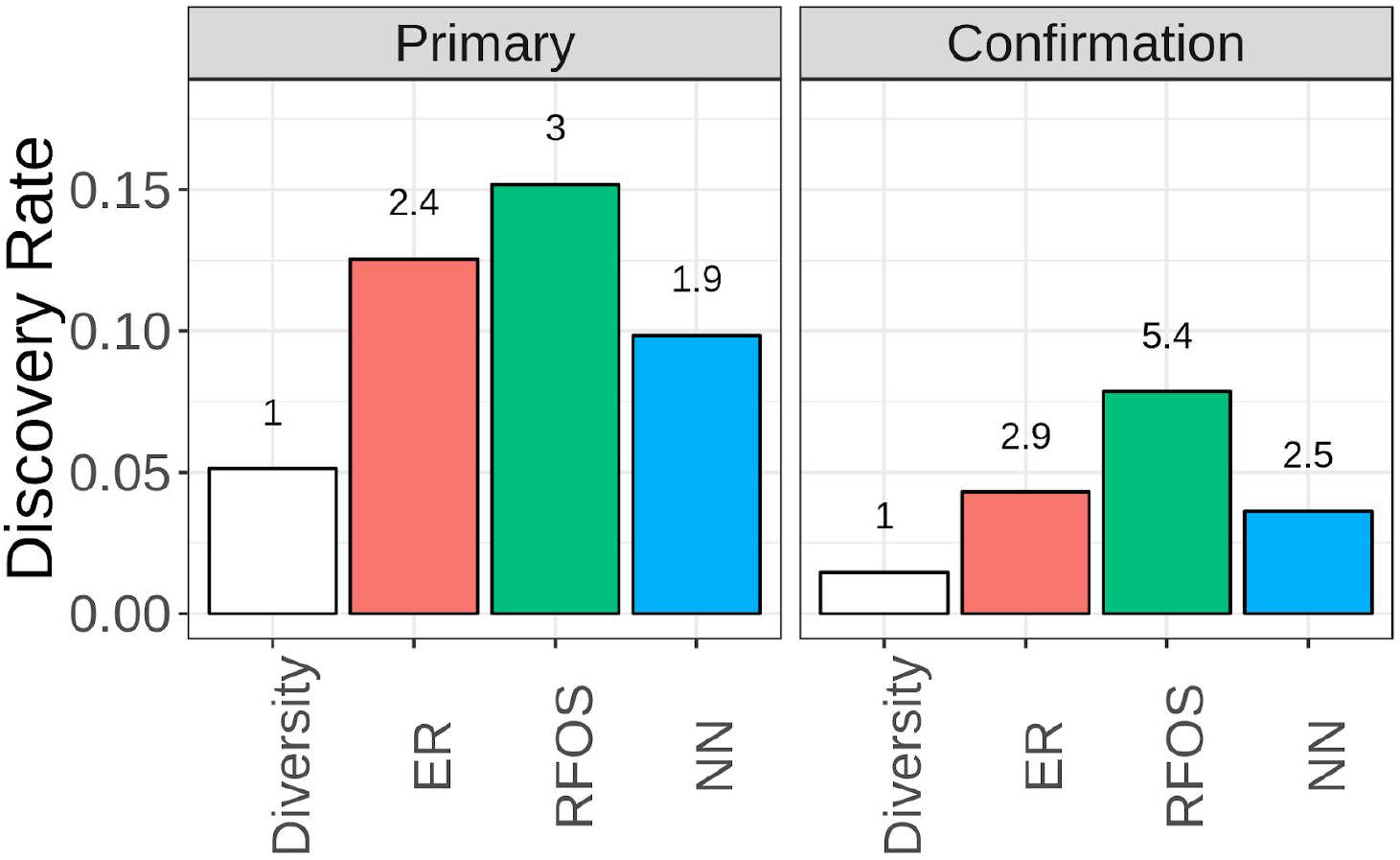
Predictive models improved discovery rate. The hit rate in the primary and confirmation screens is shown by the bars, and the fold improvement compared to the diversity screen (a random set of microbes from the collection) is displayed above each bar. RFOS improved the discovery rate by 3-fold in the primary screen, and by 5.4-fold in the confirmation screen.

The NN model did not perform as well as the RFOS model in the context of this *post-hoc* analysis. We found that the NN primary discovery rate was 1.9-fold greater than the diversity screen discovery rate (p-value of 0.01 by one-sided two-proportions Z test), and the confirmation discovery rate was 2.5-fold greater than the diversity screen (p-value of 0.03 by one-sided two-proportions Z test) (Figure 4). The precision was not statistically different from the null model, but the recall was better than the null (p-value of 0.098 compared to 1,000 random draws) (Supplemental Figure 7). Recall was not significantly better than the null model when predicted the results of the SA confirmation screen—the NN model correctly predicted two of 12 (17%) isolates that reproducibly controlled SA (p-value of 0.12, compared to 1,000 random draws). In contrast to the RFOS model, the true and false positives consisted primarily of members of the genera *Bacillus_A*, while most true negatives were classified as *Bacillus*, and most false negatives were classified as *Pseudomonas_E*.

Finally, we found that the ER primary discovery rate was 2.4-fold greater than the diversity screen discovery rate (p-value of 7.4×10^−5^ by one-sided two-proportions Z test), and the confirmation discovery rate was 2.9-fold greater than the diversity screen (p-value of 6.1×10^−3^ by one-sided two-proportions Z test) (Figure 4). The ER method outperformed the null model in terms of recall (Supplemental Figure 7). It correctly predicted 19 of the 32 (59.4%) SA-controlling isolates in the primary screen (p-value of 0.007 compared to 1,000 random draws), and three of the 12 actives against SA in the confirmation screen (p-value of 0.031 compared to 1,000 random draws). To describe model performance in taxonomic terms, most of the true and false positives were classified as *Bacillus_A*, most of the true negatives as *Bacillus*, and most of the false negatives as *Pseudomonas_E* and *Lysobacter*.

The ER method (uniquely among the three computational methods used) discovered active isolates from a novel genus which was not present in the training data—*Bacillus_I endophyticus*. These *Bacillus_I* isolates were prioritized by the enrichment method based on the presence of the BGC family “nrps2550”. The BGC family “nrps2550” shares homology with the BGC known to produce Elsinochrome A. Elsinochrome A is a perylenequinone, a family of molecules which are known to exhibit fungicidal activity by production of reactive oxygen species (29,30).

## Discussion

Many of the isolates with confirmed activity against SA and BS are from taxonomic groups that have long been known to exhibit fungicidal properties, specifically *Lysobacter (31)*, and a variety of Pseudomonads and Bacilli (Table 1) (32). Some taxa discovered in this work had not been reported to control fungal disease before, including *Paenibacillus_B sp001894745* (sometimes labeled in NCBI as *Paenibacillus alvei* or lacking species classification), MA-N2 sp002009585, a member of the family Micrococcaceae (labeled in NCBI as a member of the genus *Arthrobacter* and lacking a species identifier), a novel member of the genus *Glutamicibacter*, and a novel member of the family Planococcaceae. The broad diversity of bacterial taxa capable of controlling SA and BS suggests the widespread impact of bacterial-fungal interactions over the course of evolutionary time (33). The diversity also suggests that bacteria yet harbor many untapped opportunities for fungal control, both in agricultural and medical applications.

The gene clusters we observed to be enriched and predictive of bacterial anti-SA and anti-BS activity were a mix of homologs to well-studied BGCs and novel, uncharacterized BGCs. Many of the BGC families are highly homologous to the characterized BGCs for known fungicidal compounds (e.g. Pyrrolnitrin, Zwittermycin, Fengycin, and Surfactin). This is a strong validation of the approaches (enrichment and permutation feature importance) we used to rank BGC families, and is encouraging of future work to characterize the novel BGCs we observed. It is unclear, based on genome assemblies alone, to what extent these BGCs with homology to known BGCs produce identical, or merely related, compounds. It is also likely that some of the BGCs we found to be associated with antifungal activity are simply correlated with the taxonomic group that exhibited activity and are unrelated to the mode of action. A similar challenge is that singleton BGCs representing novel antifungal pathways cannot be detected by the approach used here. The nature of the training data make it impossible to determine which BGC families actually cause anti-SA or -BS activity. Therefore, it is necessary to do more in-depth studies to demonstrate causation, and to elucidate potential MOAs associated with BGC families discovered here.

Regardless of whether the BGCs discovered here represent the actual source of antifungal activity or are merely correlated with it, they are predictive of antifungal activity against SA and BS. Interestingly, we observed *in silico* that it is rare for BGC families to be predictive of activity outside the genus level (Figure 3C). In some cases we observed that the predictive models could correctly predict activity among isolates from a genus not represented in the training data, but *In silico*, these cases were rare and were not common enough to differentiate from random chance. It is unclear whether the genus-level limit to BGC predictive value is reflective of biology (i.e. antifungal MOAs tend to be limited to within a bacterial genus) or whether it is an artifact of the way we defined and ranked BGC families. For example, despite the *in silico* results, in the validation experiment the enrichment method did in fact lead us to discover a novel genus. The discovery of a novel genus was possibly a stroke of luck, but it is also likely that some antifungal BGC families span and are predictive across broader taxonomic groups. An example in our training data of such a BGC family is “others2810” (Table 2), which can be found (in our data set) across the class Gammaproteobacteria. In the validation experiment, the BGC family “nrps2550”—which was limited to the genus *Bacillus_A* in the training set—was the clue that led to the discovery of activity against SA by a member of the genus *Bacillus_I*. By predicting activity based on genomic features rather than whole-genome homology or taxonomic relatedness, it becomes possible to anchor predictions on features that have been broadly inherited or horizontally dispersed. In this way, genomic features act as stepping stones to new taxa with potentially novel properties for SA or BS control.

Broader and deeper data sets are needed if we wish to predict SA or BS control with more confidence. While every computational approach we tested performed significantly better than the random, null model for predicting activity in the primary assay (Figure 3B), none of the modeling approaches achieved average F1 scores greater than 20%, or maximum F1 scores greater than 40%. The relatively low model confidence highlights the complex nature of antifungal interactions, which can be influenced by subtle differences between isolates. Small changes in BGC enzymes may result in the production of different metabolites, which may or may not be antifungal. Small changes to promoters, signalling molecules, or growth conditions can all have a large impact on the observed antifungal activity against SA or BS. Furthermore, it is likely that many active isolates employed unique modes of action which could not be used to predict activity in any other isolates. All of this simply points to the need for more screening data, and the use of input features capable of capturing more subtle differences between isolates. Regardless, we found that RFOS and NN models performed the best *in silico* (Figure 3), and that predictive models improved the efficiency of our screening efforts (Figure 4). There is a lot left to learn and discover about bacterial control of SA and BS, and computational models will continue to accelerate the discovery effort.

## Acknowledgements

We express our thanks to Dr. Thomas Isakeit at Texas A&M University for providing isolates of *C. sublineolum*. We are grateful to Dr. Amos Emitati Alakonya (a banana pathologist at IITA, Nigeria) for providing the *M. fijiensis* isolate used in this study. We express our thanks to the Bill and Melinda Gates Foundation for their generous support of this work. We further thank Dr. Gregory Medlock at the University of Virginia for his thoughtful comments and suggestions during the preparation of this manuscript. And none of this work would have been possible without the phenomenal team at AgBiome!

## Creative Commons Attribution

This work was supported, in whole or in part, by the Bill & Melinda Gates Foundation [OPP1151824]. Under the grant conditions of the Foundation, a Creative Commons Attribution 4.0 Generic License has already been assigned to the Author Accepted Manuscript version that might arise from this submission.

