## Supplemental Information for "Genomics-accelerated discovery of diverse fungicidal bacteria"

**Supplemental Materials for “Genomics-accelerated discovery of diverse fungicidal bacteria”**

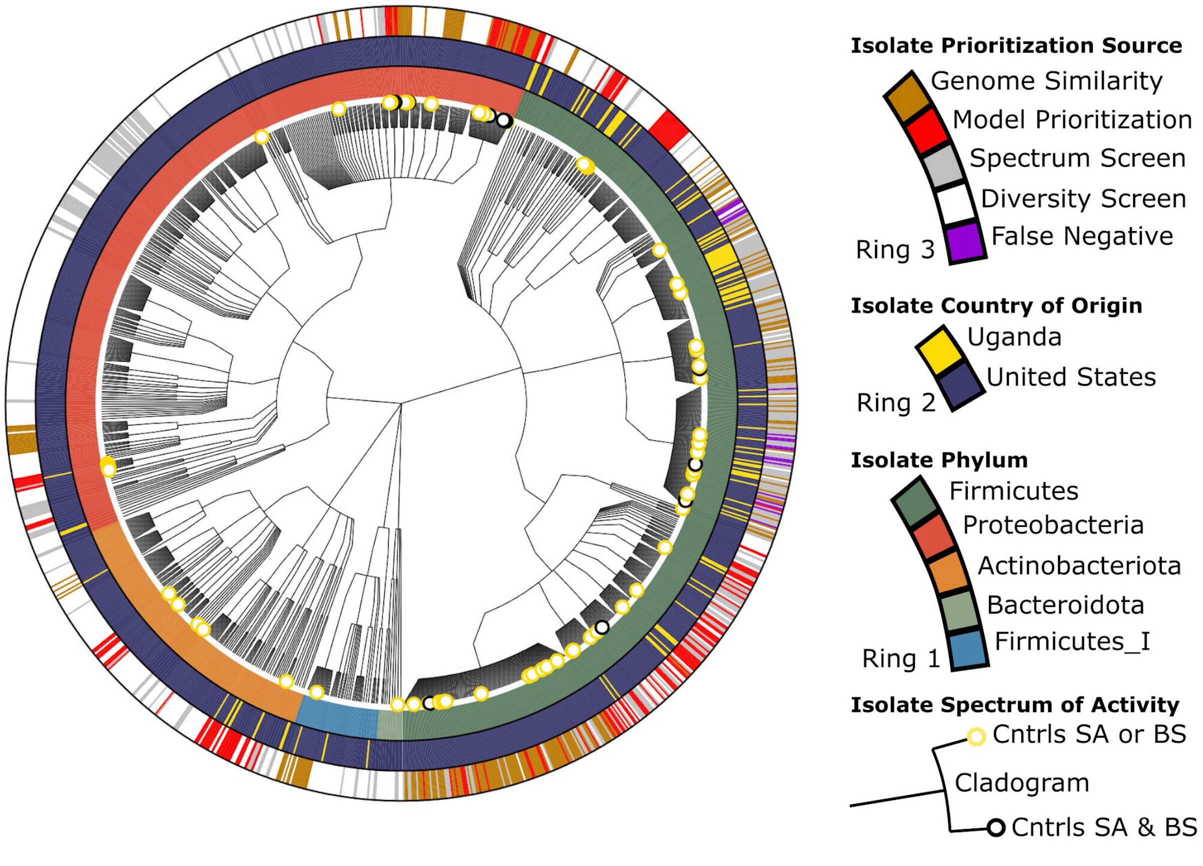


**Supplemental Figure 1. Full collection of strains screened against SA and BS.** The cladogram in the center highlights the diversity of the biocontrol strains discovered in this work (five phyla represented by the center branches, 14 genera, 22 species). Multiple isolates displayed activity against both diseases (black leaf nodes). The outer ring shows the isolate prioritization strategies that led to each discovery, including predictive models. The “false negative” isolates in purple were predicted to be inactive by all three modeling approaches and yet were reproducibly active against SA.

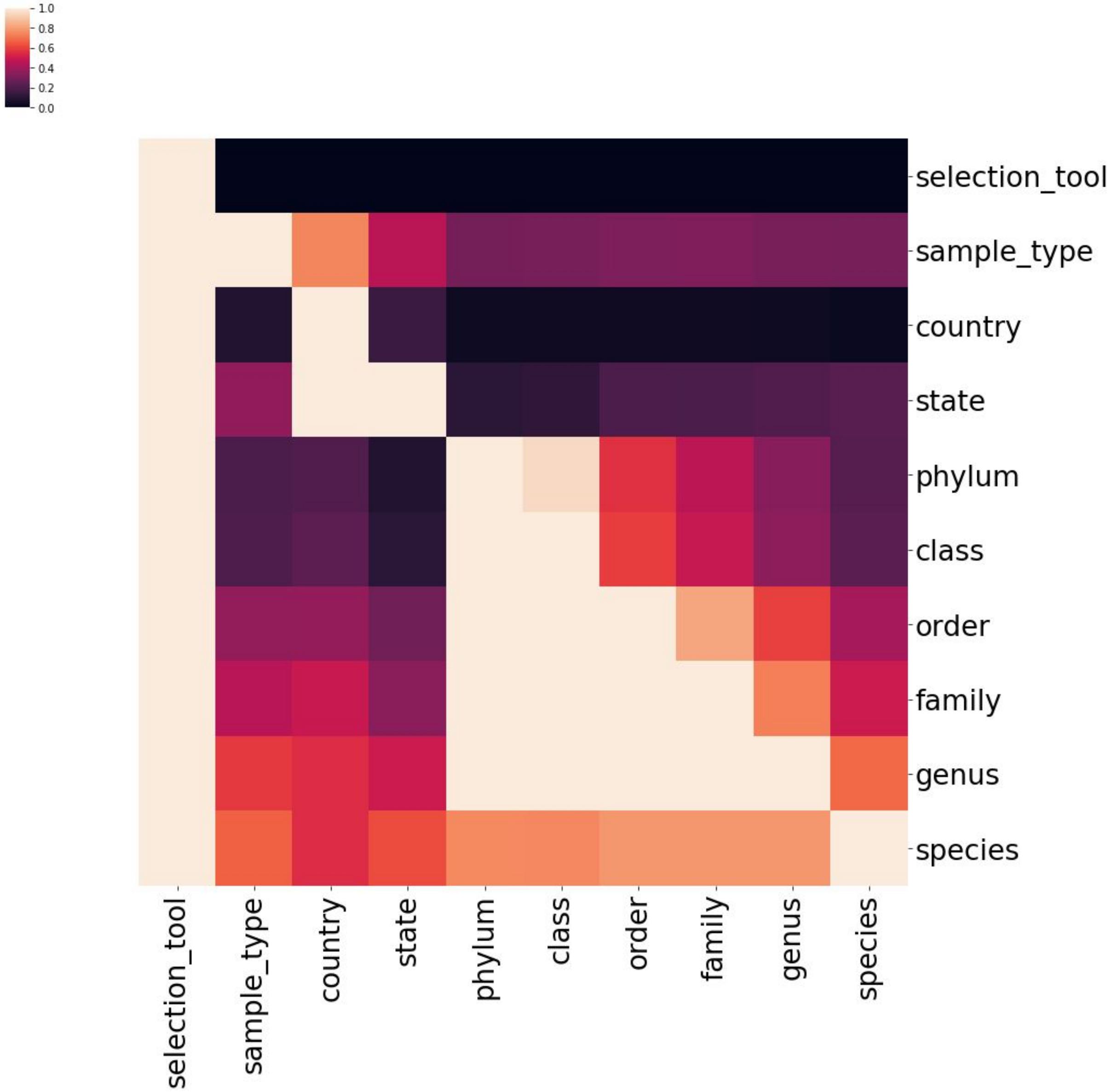


**Supplemental Figure 2**. **Covariance between metadata categories.** Theil's U is the uncertainty of x given y, where an output close to 0 means y provides no information about x, and a value close to 1 means y provides full information about x. You read the figure as "if you know the value from the row, how much information does that provide about the value in the column?" For example, if you know the **genus**, it gives you full information about the **family**, **order**, **class** and **phylum**, but the opposite is not true.

| Supplemental Table 1 | | | | | |
|---|---|---|---|---|---|
| **BGC Family ID** | **99th Percentile Importance** | **Top Enrichment Scores** | **Predicted Product Class** | **Homology to Known BGC** | **Taxonomic Distribution** |
| nrps4050 | TRUE | TRUE | NRPS | Bacillibactin | genus:Bacillus_A |
| nrps4047 | TRUE | TRUE | NRPS;T1PKS | Zwittermycin A | genus:Bacillus_A |
| others9203 | TRUE | TRUE | siderophore | Petrobactin | genus:Bacillus_A |
| nrps3014 | TRUE | FALSE | NRPS | None | species:Bacillus_A toyonensis |
| others7677 | TRUE | FALSE | other | Bacilysin | genus:Bacillus |
| ripps2504 | TRUE | FALSE | LAP;bacteriocin | None | species:Bacillus_A toyonensis |
| others3454 | TRUE | FALSE | betalactone | None | genus:Bacillus |
| ripps7679 | TRUE | FALSE | bacteriocin | None | genus:Bacillus |
| nrps7678 | TRUE | FALSE | NRPS | Surfactin | genus:Bacillus |
| pksother12634 | TRUE | FALSE | NRPS;transAT-PKS;betalactone | Fengycin | species:Bacillus velezensis |
| ripps11067 | TRUE | FALSE | LAP | Plantazolicin | species:Bacillus safensis |
| terpene7676 | TRUE | FALSE | siderophore | Carotenoid | genus:Bacillus |
| ripps3993 | TRUE | FALSE | bacteriocin | None | genus:Bacillus_A |
| ripps10679 | TRUE | FALSE | bacteriocin | None | species:Bacillus safensis |
| nrps1603 | TRUE | FALSE | NRPS | Pyoverdine | species:Pseudomonas_E extremorientalis |
| terpene1433 | TRUE | FALSE | terpene | Carotenoid | species:gtdb_novel_strain |
| others4222 | TRUE | FALSE | CDPS | None | genus:Pseudomonas_E |
| ripps2941 | TRUE | FALSE | bacteriocin | None | genus:Pseudomonas_E |
| nrps4841 | TRUE | FALSE | NRPS | Syringomycin | species:Pseudomonas_E protegens |
| nrps4795 | FALSE | TRUE | NRPS | Pyoverdine | species:Pseudomonas_E protegens |
| nrps2943 | FALSE | TRUE | NRPS | Lipopeptide 8D1-1/2 | species:Pseudomonas_E protegens |
| others2810 | FALSE | TRUE | other | Pyrrolnitrin | class:Gammaproteobacteria |
| pksother4889 | FALSE | TRUE | PKS-like | Anikasin | species:Pseudomonas_E protegens |
| others3032 | FALSE | TRUE | arylpolyene | APE Vf | genus:Pseudomonas_E |
| ripps4856 | FALSE | TRUE | bacteriocin | None | genus:Pseudomonas_E |
| pksother2939 | FALSE | TRUE | T3PKS | 2,4-Diacetylphloroglucinol | family:Pseudomonadaceae |
| nrps2808 | FALSE | TRUE | NRPS | Thiazostatin | genus:Pseudomonas_E |
| others3030 | FALSE | TRUE | NAGGN | None | family:Pseudomonadaceae |

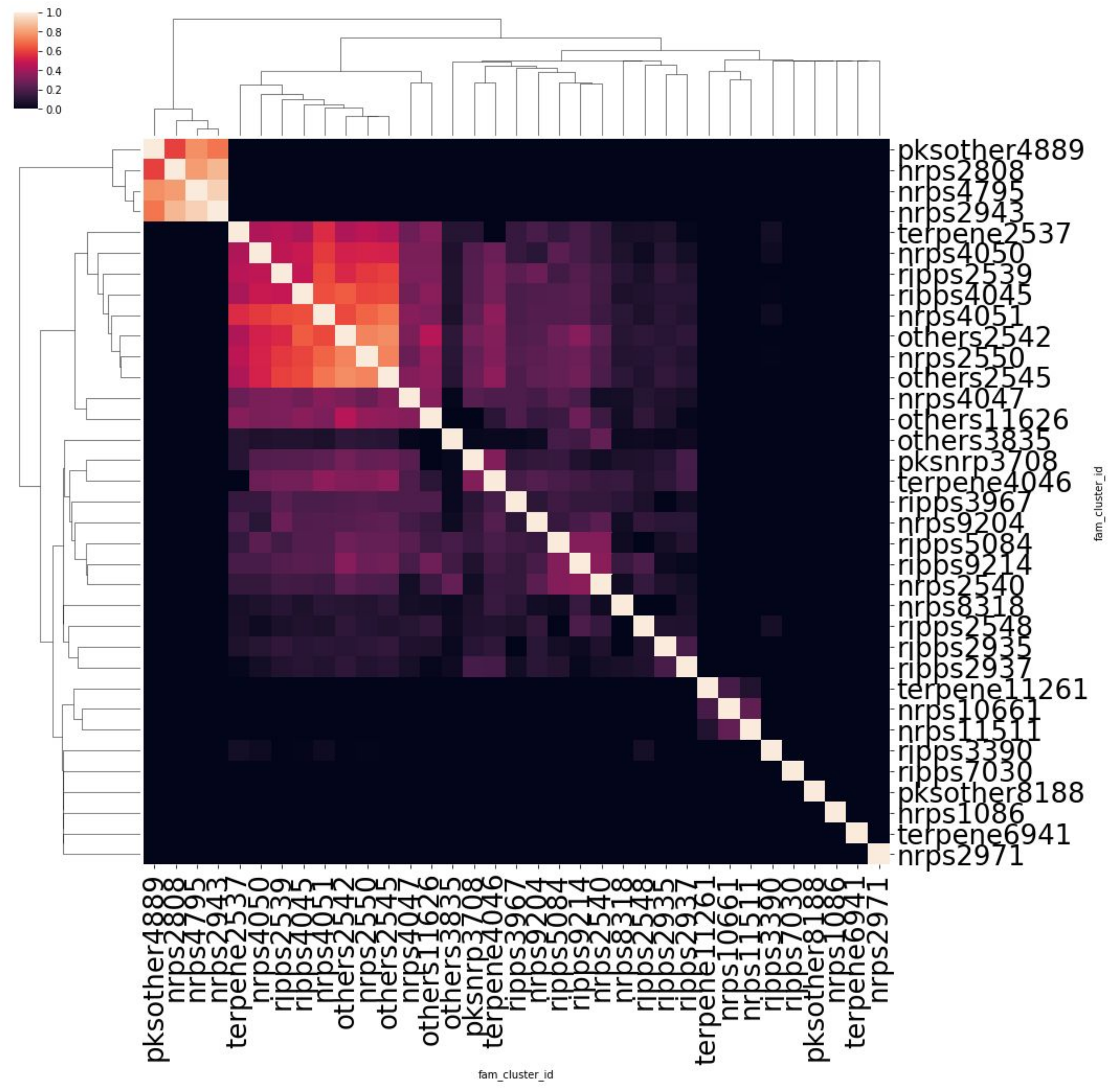


**Supplemental Figure 3**. **SA features co-occurrence, calculated as the Jaccard similarity between the sets of isolate genomes that contain members of the BGC families.** For example, “pksother4889” was not found in any of the same genomes as “nrps7030”, and so the Jaccard similarity is 0. On the other hand, “terpene2537” was found in many of the same genomes as “nrps4051”.

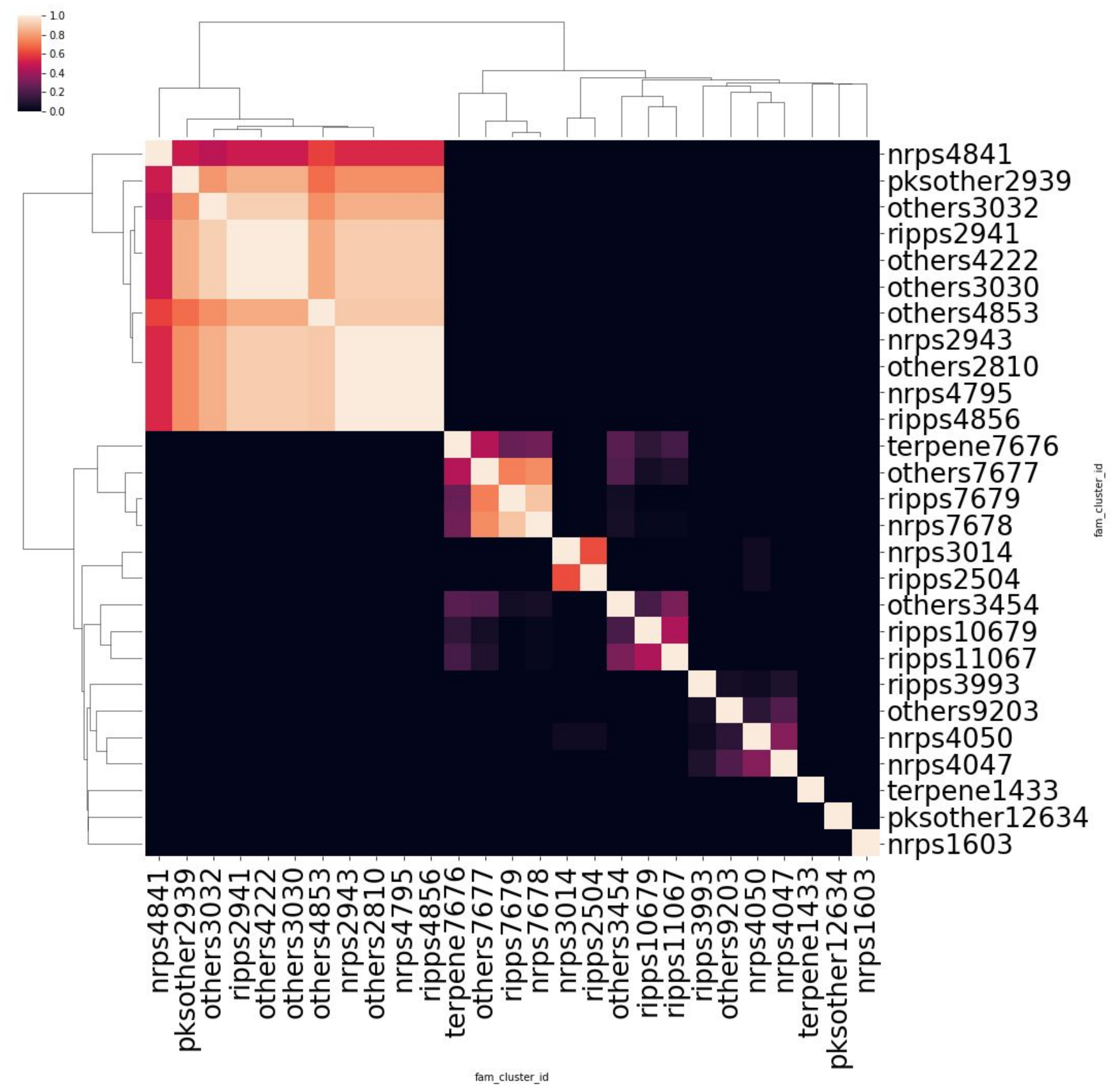


**Supplemental Figure 4. BS features co-occurrence, calculated as the Jaccard similarity between the sets of isolate genomes that contain members of the BGC families.** For example, “nrps4841” was not found in any of the same genomes as “pksother12634”, and so the Jaccard similarity is 0. On the other hand, “nrps4841” was found in many of the same genomes as “pksother2939”.

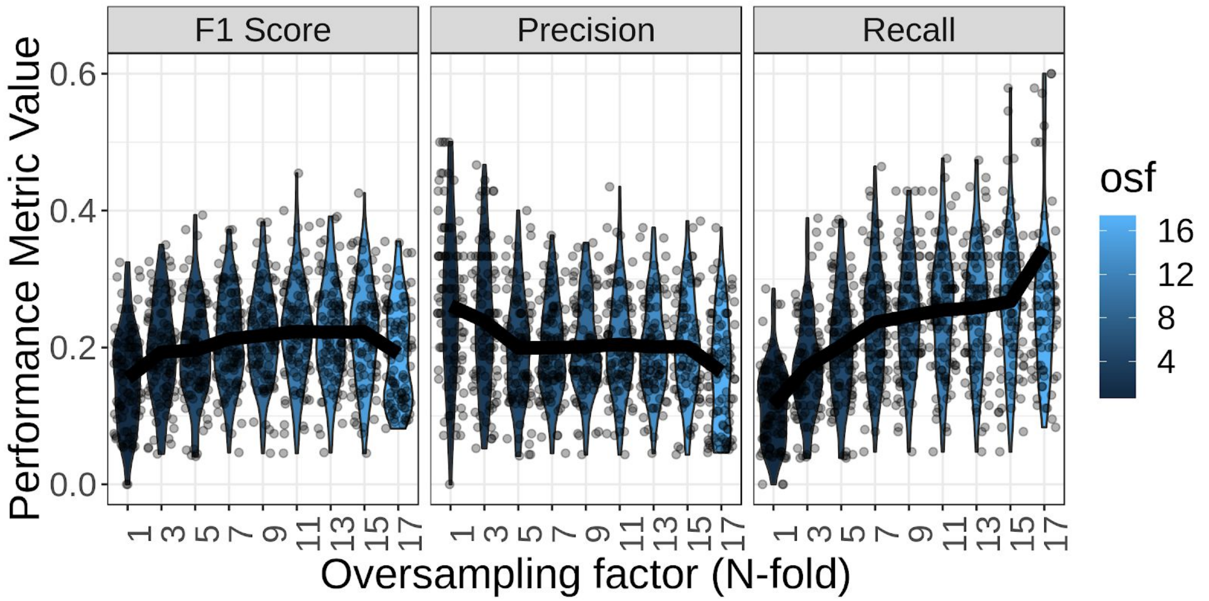


**Supplemental Figure 5. The effect on random forest model performance of oversampling positive observations in the SA primary screen data set.**

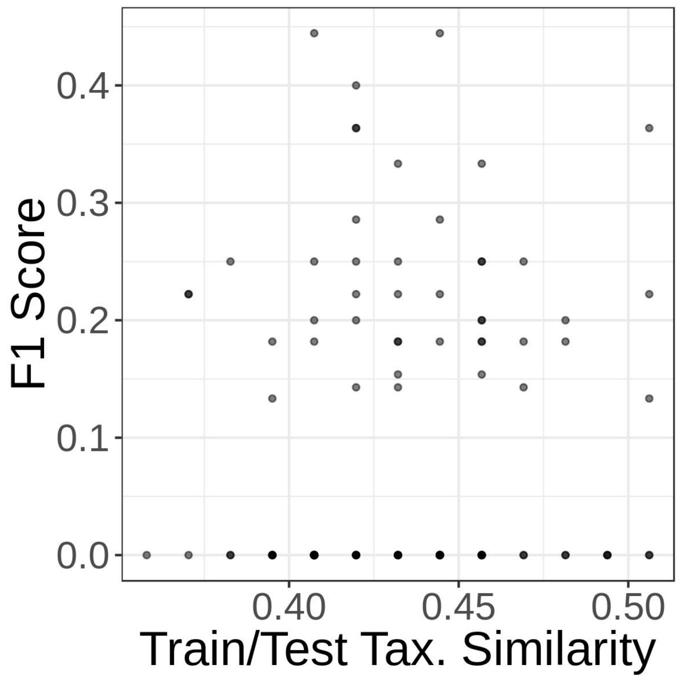


**Supplemental Figure 6. Correlation between the taxonomic overlap of the training/test set and the model performance.**

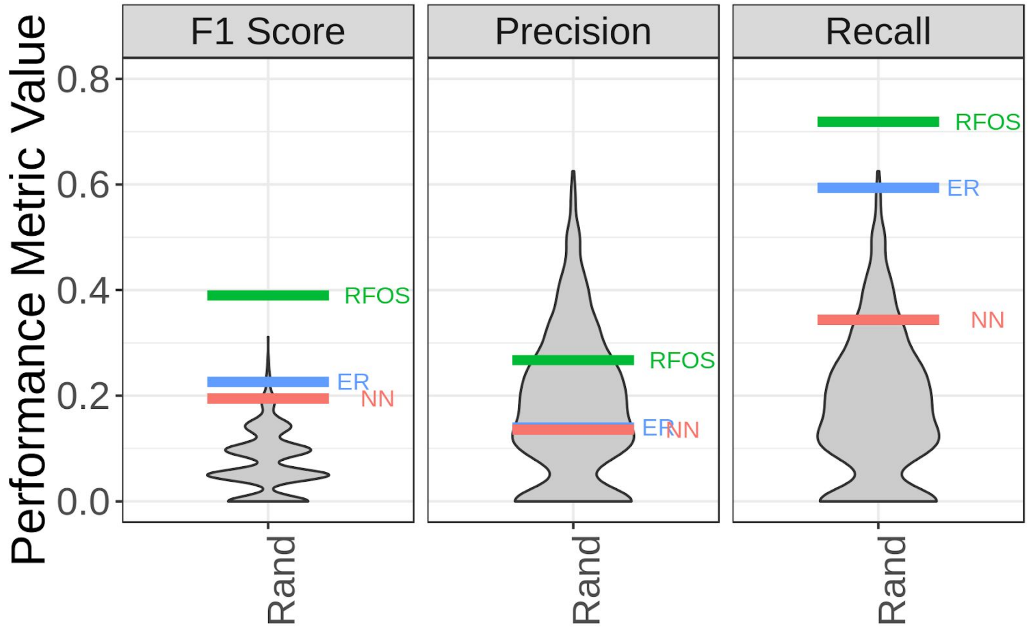


**Supplemental Figure 7. Performance of predictive models during *in vivo* validation experiment with 176 isolates.** The gray distribution represents the results of 1,000 draws from a random, null model. The horizontal lines indicate how each predictive model performed on this single validation set of 176 isolates. The closer to the upper tail of the null distribution, the more valuable the performance. The closer to the mean, the less different from random chance. In this case, the models tended to achieve better recall (driving up the F1 score) than random chance would suggest.

| **Supplemental Table 2** | |
|---|---|
| **BGC Family ID** | **Feature Selected By** |
| ripps7574 | random_forest |
| nrps4051 | enrichment |
| nrps9204 | random_forest |
| others3835 | random_forest |
| nrps2567 | random_forest |
| terpene7676 | random_forest |
| nrps4050 | enrichment |
| nrps8190 | random_forest |
| ripps3390 | both |

| | |
|---|---|
| nrps2597 | random_forest |
| ripps2539 | both |
| others7673 | random_forest |
| nrps4071 | random_forest |
| nrps2936 | random_forest |
| others2545 | both |
| nrps2969 | random_forest |
| nrps10526 | random_forest |
| nrps8318 | random_forest |
| others9203 | random_forest |
| ripps2548 | random_forest |
| pksother4889 | enrichment |
| nrps9044 | random_forest |
| ripps3967 | random_forest |
| others4222 | random_forest |
| nrps6166 | random_forest |
| nrps4047 | random_forest |
| ripps10679 | random_forest |
| ripps3960 | random_forest |
| ripps4045 | enrichment |
| nrps2550 | enrichment |
| nrps2943 | both |
| nrps4795 | enrichment |
